# Methylene Blue has a potent antiviral activity against SARS-CoV-2 in the absence of UV-activation in vitro

**DOI:** 10.1101/2020.08.14.251090

**Authors:** Valeria Cagno, Chiara Medaglia, Andreas Cerny, Thomas Cerny, Caroline Tapparel, Erich Cerny

## Abstract

Methylene blue is an FDA and EMA approved drug with an excellent safety profile. It displays broad-spectrum virucidal activity in the presence of UV light and has been shown to be effective in inactivating various viruses in blood products prior to transfusions. In addition, its use has been validated for methemoglobinemia and malaria treatment. Here we show the virucidal activity of methylene blue at low micromolar concentrations and in the absence of UV activation against SARS-CoV2.

## INTRODUCTION

Viral pandemics cause significant morbidity and mortality. Influenza A viruses and coronaviruses are among the major human threats due to the presence of an important animal reservoir and the virus ability to cross the species barrier via mutation and recombination. The ongoing SARS-CoV2 pandemic unveiled the inability to develop specific antivirals and vaccines in a short time frame and the need for broad-spectrum antiviral drugs ready to use when a new pandemic begins.

Methylene Blue (MB) is an FDA and EMA approved drug with an excellent safety profile. Due to its antimicrobial, anti-inflammatory and antitoxic effects, photoactivated MB is used for a wide range of applications including treatment of methemoglobinemia or malaria (reviewed in (1)). Photoactivated MB is also widely used to obtain blood product preparations free of viruses such as HIV (2), Ebola or Middle East Respiratory Syndrome coronavirus (MERS) (3), or SARS-CoV2 virus (4). The antiviral activity appears to rely on multiple mechanisms, and is more potent for enveloped viruses (5). Among other actions, MB is known to corrupt DNA or RNA integrity. This action is based on a redox reaction in which the molecule takes electrons on its aromatic thiazine ring, is reduced to leuko-methylene blue (MBH_2_) and transfers electrons to other compounds. MB as a sensitizer in combination with oxygen and a source of energy results in the production of singlet oxygen, a very reactive reaction partner which induces guanine oxidation (8-oxo-7,8-dihydroguanine (8-oxoGua) lesions) damaging DNA or RNA. Other mechanisms include but are not limited to a) modified carbonyl moieties on proteins, b) single-strand breaks (ssb) in the RNA genome c) RNA-protein crosslinks, all lesions correlating well with a broad-spectrum virucidal activity (6).

MB presents few side effects at doses below 5mg/kg. Long term, high dose treatment such as for malaria may result in temporary fully reversible blueish coloration of urine, sclera and skin. Sensitivity to thiazine dyes and glucose 6 phospate dehydrogenase (G-6-PD) deficiency are contraindications (7).

Using a classical virus neutralization assay for SARS-CoV-2 we present in vitro data demonstrating a potent antiviral activity without UV-activation, opening the possibility to evaluate MB for the treatment of patients. The strong antiviral activity without light (experiments performed in a closed stainless-steel box) was evidenced with 20 hours incubation time.

## RESULTS AND DISCUSSION

Different doses of MB were incubated with 5*10^6^ plaque forming units (pfu) of SARS-CoV2 for 20 hours in the dark at room temperature. The 50% cytotoxic concentration (CC_50_) of MB was determined to be 84.72 μg/ml and only lower doses were evaluated for efficacy. The results shown in Figure 1 evidence a complete inhibition up to 10 µg/ml and a 3.05 log reduction at 0.08 µg/ml, the lowest dose tested. Of note Jin and colleagues (4) observe no virucidal activity of MB against SARS-CoV2 when treatment is run for 20’ in the dark. Additional experiments with shorter incubation times are underway to determine the minimal exposure time necessary for activity in the dark.

**Figure 1.**
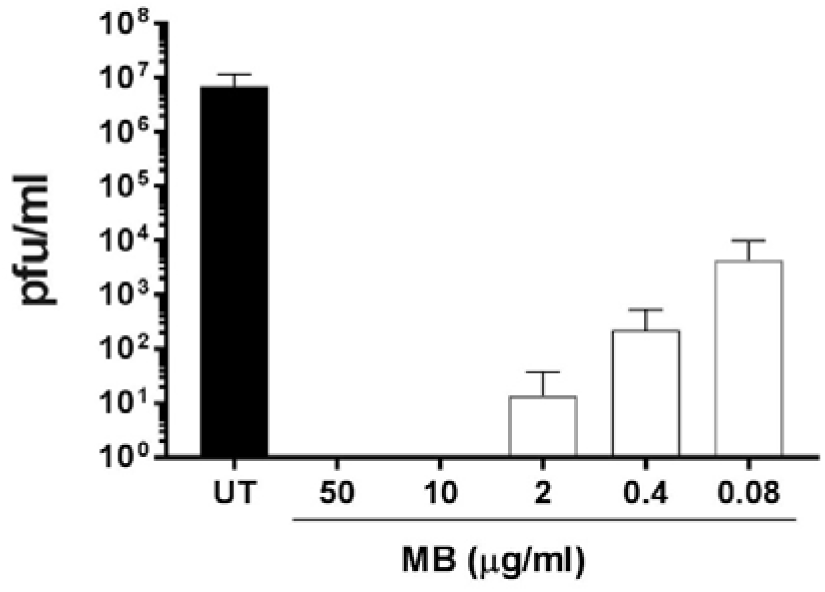
SARS CoV-2 virucidal assay. MB was incubated with SARS-CoV2 (5*10^6^ pfu) for 20h in the dark. At the end of the incubation, mixtures were serially diluted and added for 1h at 37°C on Vero-E6 cells. Mixtures were then removed and cells overlaid with medium containing 0.8% avicel. Cells were fixed 48hpi and plaques were counted in order to determine the viral titer in presence or absence of MB. Results are mean and SD of three independent experiments. 50 µg/ml (156 µM), 10 µg/ml (31.2 µM), 2 µg/ml (6.25 µM), 0.4 µg/ml (1.25 µM), 0.08 µg/ml (0.25 µM).

Subsequently we evaluated the possible combinatorial activity of MB and immunoglobulins present in the serum of a convalescent individual. Preliminary tests were conducted to identify the optimal dose of serum to use and evidenced a 4.15 log decrease in presence of 1:16 dilution, a 2.3 log decrease in presence of 1:80 dilution, and absence of neutralization at lower serum dilutions (data not shown). In order to be able to evaluate the additive effects, the serum was diluted 1:80 for further experiments. Incubations for 20 hours in the dark were then carried out with SARS-CoV2, three different doses of MB in presence or absence of diluted convalescent serum. The results (Figure 2) evidence a partial additive effect of MB and convalescent sera, with a complete neutralization with the combination of MB and serum up to 0.4 µg/ml of MB, supporting both the use of MB combined to convalescent serum to treat patients in vivo and the beneficial effect of MB treatment in patients having started to mount an antibody response.

**Figure 2.**
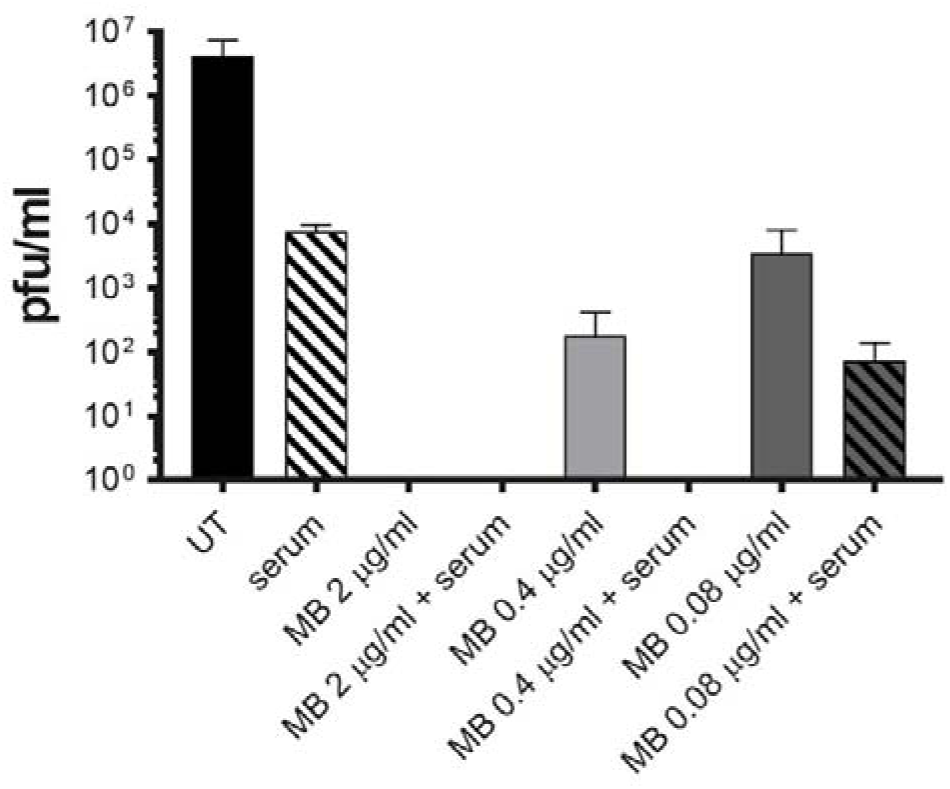
SARS CoV-2 virus neutralization assay in presence of convalescent serum. Methylene Blue (MB) alone or in combination with 1:80 diluted human convalescent serum was incubated with SARS-CoV2 (5*10^6^ pfu) for 20h in the dark. At the end of the incubation mixtures were serially diluted and added for 1h at 37°C on Vero-E6 cells. Mixtures were then removed and cells overlaid with medium containing 0.8% avicel. Cells were fixed 48hpi and plaques were counted in order to determine the viral titer in presence or absence of MB and serum. Results are mean and SD of three independent experiments.

Altogether, our results support the possibility to use MB in clinical studies against SARS-CoV2 in the initial phases of the disease. Based on its known mechanisms of actions, MB will destroy extracellular viruses limiting viral replication and spread beyond the upper respiratory tract. In addition, its anti-inflammatory activity will lower the side effects linked to the host response. The effect of MB on intracellular SARS-CoV2 needs to be tested as well as its efficacy when administered after infection start. This will help to find out if MB also presents a therapeutic interest when administered at more advanced disease stages.

## MATERIALS AND METHODS

### Compound and convalescent serum

Methylene blue (Methylthonium chloride solution) was purchased from ProVepharm. A convalescent serum tested positive for IgG against SARS-CoV2 by ELISA was separated through centrifugation, aliquoted and stored at −80°C.

### Cells and Virus

Vero C1008 (clone E6) (ATCC CRL-1586) were a kind gift from Prof Gary Kobinger, and were propagated in DMEM High Glucose + Glutamax supplemented with 10% fetal bovine serum (FBS) and 1% penicillin/streptavidin (pen/strep).

SARS-CoV2/Switzerland/GE9586/2020 was isolated from a clinical specimen in the University Hospital of Geneva in Vero-E6. Cells were infected and supernatant was collected 3 days post infection, clarified, aliquoted and frozen at −80°C before titration by plaque assay in Vero-E6.

### Toxicity assay

Vero-E6 (13000 cells per well) were seeded in 96-well plate. Methylene blue was serially diluted in DMEM supplemented with 5% FBS and added on cells for 1h, followed by a washout, addition of DMEM supplemented with 5% FBS for additional 48h hours. MTS reagent (Promega) was added on cells for 3h at 37°C according to manufacturer instructions, subsequently absorbance read at 570 nm. Percentages of viability were calculated by comparing the absorbance in treated wells and untreated. 50% cytotoxic concentration (CC_50_) were calculated with Prism 8 (GraphPad, USA)

### Neutralization assay

SARS-CoV2 5*10^6^ pfu was incubated at room temperature in the dark (in an opaque stainless steel box) in a final volume of 400 µl with different dilutions of MB and or of MB plus convalescent serum for 20h. At the end of the infection mixtures were serially diluted (1:10 factor) and added on Vero cells (100’000 cells per well) seeded in 24-well plate 24h in advance, for 1h at 37°C. The monolayers were then washed and overlaid with 0.8% avicel rc581 in medium supplemented with 5% FBS. Two days after infection, cells were fixed with paraformaldehyde 4% and stained with crystal violet solution containing ethanol. Plaques were counted, and the titer of the different conditions was calculated. Statistical analysis was done with Prism software (Prism 8, GraphPad).

## AKNOWLEDGEMENTS

E. Cerny, Th. Cerny, A. Cerny, C. Tapparel have a Swiss patent application pending covering the use of Methylene blue as an anti-viral drug for SARS-CoV-2. E. Cerny is owner of Omni Drugs SA, which has no ties to the present project. The authors declare no conflicting interests concerning the present subject matter.

